# Antibody-mediated depletion of select T cell subsets in blood and tissue of nonhuman primates

**DOI:** 10.1101/2023.12.22.572898

**Authors:** Matthew S. Sutton, Allison N. Bucsan, Chelsea C. Lehman, Megha Kamath, Supriya Pokkali, Diogo M. Magnani, Robert Seder, Patricia A. Darrah, Mario Roederer

**Affiliations:** Vaccine Research Center, National Institute of Allergy and Infectious Diseases (NIAID), National Institutes of Health (NIH), Bethesda, MD, USA; Nonhuman Primate Reagent Resource, University of Massachusetts Chan Medical School, Worcester, MA, USA

**Keywords:** *in vivo* depletion, nonhuman primates, MT807R1, CD4R1, CD8β255R1, C207, tissue leukocytes

## Abstract

Understanding the immunological control of pathogens requires a detailed evaluation of the mechanistic contributions of individual cell types within the immune system. While knockout mouse models that lack certain cell types have been used to help define the role of those cells, the biological and physiological characteristics of mice do not necessarily recapitulate that of a human. To overcome some of these differences, studies often look towards nonhuman primates (NHPs) due to their close phylogenetic relationship to humans.

To evaluate the immunological role of select cell types, the NHP model provides distinct advantages since NHP more closely mirror the disease manifestations and immunological characteristics of humans. However, many of the experimental manipulations routinely used in mice (e.g., gene knock-out) cannot be used with the NHP model. As an alternative, the *in vivo* infusion of monoclonal antibodies that target surface proteins on specific cells to either functionally inhibit or deplete cells can be a useful tool. Such depleting antibodies have been used in NHP studies to address immunological mechanisms of action. In these studies, the extent of depletion has generally been reported for blood, but not thoroughly assessed in tissues.

Here, we evaluated four depleting regimens that primarily target T cells in NHP: anti-CD4, anti-CD8α, anti-CD8β, and immunotoxin-conjugated anti-CD3. We evaluated these treatments in healthy unvaccinated and IV BCG-vaccinated NHP to measure the extent that vaccine-elicited T cells – which may be activated, increased in number, or resident in specific tissues – are depleted compared to resting populations in unvaccinated NHPs. We report quantitative measurements of *in vivo* depletion at multiple tissue sites providing insight into the range of cell types depleted by a given mAb. While we found substantial depletion of target cell types in blood and tissue of many animals, residual cells remained, often residing within tissue. Notably, we find that animal-to-animal variation is substantial and consequently studies that use these reagents should be powered accordingly.

## 1 Introduction

The use of monoclonal antibodies (mAbs) for the study and treatment of diseases is well recognized. mAbs can also be an effective tool in mechanistic studies to acutely deplete specific cell types *in vivo* in the absence of knock-out animal models. For example, antibody-mediated depletion of CD8+ T cells in nonhuman primates (NHPs) infected with simian immunodeficiency virus (SIV) highlighted the importance of CD8+ T cells in controlling viral replication (1–4). More recently, a similar approach demonstrated the importance of vaccine-elicited CD8+ T cells in controlling replication of SARS-CoV-2 in a NHP model (5). In the clinic, mAbs that bind surface proteins on B cells, such as CD20, have been shown to be effective in the treatment of B cell lymphomas and autoimmune diseases (6–8). Targeting the surface marker of T cells with depleting mAbs specific to CD3 has also proved helpful in reducing graft-versus-host disease in transplant patients (9). Thus, the administration of mAbs has been an effective strategy for *in vivo* depletion of specific cell types in both research and clinical settings.

Unlike broad spectrum treatment approaches (such as chemotherapy) that exert their effects over a wide range of cell types, mAb specificity and affinity for only the molecule against which they were generated allows a more focused approach. When bound to its target, conventional mAbs can impact cells in multiple ways: they can alter downstream signaling pathways, directly induce apoptosis, or deplete cells through multiple Fc-mediated mechanisms (10). Some examples of these Fc-mediated mechanisms include elimination of a mAb-bound target cell through antibody-dependent cell-mediated cytotoxicity (ADCC), antibody-dependent cellular phagocytosis (ADCP), or complement-dependent cytotoxicity (CDC). Another approach for achieving *in vivo* depletion of cells expressing a target is to chemically conjugate a mAb, or its fragments, to an immunotoxin such as diphtheria toxin (11). By doing so, the toxin-conjugated mAb retains its specificity for its target and once endocytosed, the enzymatic fragment of the toxin is translocated to the cytoplasm and inhibits protein synthesis, effectively killing the cell (12, 13). Regardless of how mAbs achieve *in vivo* depletion, they provide a unique approach to better understand disease pathogenesis and evaluate new treatment regimens.

NHPs provide an invaluable resource for studying disease pathogenesis and determining immune-mediated mechanisms relevant to humans (e.g. of protection following vaccination). For antibody-mediated depletion studies, the ability to extensively sample NHP tissues allows a comprehensive assessment of the extent and location by which *in vivo* depletion occurs. Multiple mAbs exist to deplete NHP lymphocytes *in vivo*, such as ones targeting the cell surface proteins CD3, CD4, or CD8. As CD8 is expressed on T and NK cells as either a CD8αα homodimer or CD8αβ heterodimer it is possible to target cells expressing either form by using an anti-CD8α mAb, or those expressing CD8αβ by using an anti-CD8β mAb (14, 15). Targeting CD8β is thought to selectively deplete conventional CD8+ T cells, the vast majority of which express CD8αβ, without depleting donor-unrestricted T cells (MAIT, γδ) or NK cell populations that are thought to express CD8αα but not CD8αβ (16, 17). However, the delineation of CD8α and CD8β expression on conventional and donor-unrestricted cell subsets is not clear cut, complicating interpretations of CD8 depletion (14, 18–21).

Vaccination elicits antigen-specific populations that may be activated, increased in number, or compartmentalized in anatomical sites and thus may be more or less vulnerable to depleting antibodies compared to resting or naïve populations. Previously, we showed that NHP immunized intravenously (IV) with BCG exhibit a large increase in activated lymphocyte numbers in bronchoalveolar lavage (BAL) and lung tissue and are subsequently protected from *Mycobacterium tuberculosis* (*Mtb*) challenge (22). These correlative studies implicate the importance of antigen-specific CD4+ T cells in BAL, but mechanistic assessment requires depletion studies to define the necessity of these cells. In addition to CD4+ T cells, CD8+ T cells, γδ+ T cells, and MAIT cells have also been implicated in the control of tuberculosis (23–25). The differential expression of CD8α and CD8β on these cell types highlights the importance of distinguishing the isoforms on these cells and for selectively depleting them.

As *in vivo* depletion of individual lymphocyte populations could provide insight on the mechanistic correlates of IV BCG mediated protection, we first sought to determine the range and efficacy of *in vivo* depletion following immunization. We assessed depletion longitudinally in peripheral blood mononuclear cells (PBMCs), BAL, axillary lymph nodes (LNs), and bone marrow (BM) and at necropsy in spleen, lung lobes, and additional LNs within the periphery (mesenteric and inguinal) and lung (hilar and carinal). To best interpret such *in vivo* depletion studies, it is important to consider not only the extent by which depletion occurs, but where depletion occurs and what cell types may be affected. It is also critical when measuring cell depletion to provide not only representational data, such as changes to population composition (e.g. percentages), but also quantitative information about changes to the absolute cell count. This is critical because homeostatic adjustments that occur in the context of *in vivo* depletion may result in an increase in the fraction of a cell population when total cell numbers may actually be decreasing. To address these questions, we performed a comparative study with unvaccinated and IV-BCG vaccinated NHP that received one of the following: anti-CD8α depleting antibody (clone MT807R1), anti-CD8β depleting antibody (clone CD8β255R1), anti-CD4 depleting antibody (clone CD4R1), or anti-CD3 diphtheria toxin-conjugated antibody (clone C207). IV BCG vaccination was chosen as a model as it induces a large influx of T cells into the NHP BAL and lung that provide protection against *Mtb* challenge.

## Materials and Methods

### Ethics Statement

The animal protocols and procedures in this study were reviewed and approved by the Animal Care and Use Committee (ACUC) of both the Vaccine Research Center (in accordance to all the NIH policy and guidelines) as well as the Institutional Animal Care and Use Committee (IACUC) of Bioqual, Inc. where non-human primates were housed for the duration of the study. Bioqual Inc., and the NIH are both accredited by the Association for Assessment and Accreditation of Laboratory Animal Care (AAALACi) and are in full compliance with the Animal Welfare Act and Public Health Service Policy on Humane Care and Use of Laboratory Animals. In accordance to the institutional policies of both institutions, all compatible non-human primates are always pair-housed, and single housing is only permissible when scientifically justified or for veterinary medical reasons, and for the shortest duration possible.

### BCG vaccination

Male and female rhesus macaques were vaccinated under sedation in two cohorts. BCG Danish Strain 1331 (Statens Serum Institut, Copenhagen, Denmark) was expanded (26), frozen at approximately 3 × 10^8^ colony forming units (CFUs) ml^−1^ in single-use aliquots and stored at −80 °C. Immediately before injection, BCG was thawed and diluted in cold PBS containing 0.05% tyloxapol (Sigma-Aldrich) and 0.002% antifoam Y-30 (Sigma-Aldrich) to prevent clumping and foaming. (27). IV BCG (5 × 10^7^ CFUs) was injected into the left saphenous vein in a volume of 2 mL. Text refers to nominal BCG doses—actual BCG CFUs were quantified immediately after vaccination and were calculated at 2.0 x 10^7^ CFUs or 4.8 x 10^7^ CFUs depending on the cohort.

### Rhesus blood, BAL, and tissue processing

Blood PBMCs were isolated using Ficoll-Paque PLUS gradient separation (GE Healthcare Biosciences) and standard procedures; BAL wash fluid (3 × 20 ml washes of PBS) was centrifuged and cells were combined before counting, as described (28). LNs were mechanically disrupted and filtered through a 70-μm cell strainer. Bone marrow was filtered through a 70-μm cell strainer and further separated using Ficoll-Paque. Lung and spleen tissues were processed using gentleMACS C Tubes and Dissociator in RPMI 1640 (ThermoFisher Scientific). Spleen mononuclear cells were further separated using Ficoll-Paque. Lung tissue was enzymatically digested using collagenase (ThermoFisher Scientific) and DNase (Sigma-Aldrich) for 30–45 minutes at 37 °C with shaking, followed by passing through a cell strainer. Single-cell suspensions were resuspended in warm R10 (RPMI 1640 with 2 mM L-glutamine, 100 U ml^−1^ penicillin, 100 μg ml^−1^ streptomycin, and 10% heat-inactivated FBS; Atlantic Biologicals) or cryopreserved in CryoStor (BioLife Solutions, Inc.) or FBS containing 10% DMSO in liquid nitrogen.

### Administration of *in vivo* depleting mAb

The NIH Nonhuman Primate Reagent Resource (NNHPRR) engineered and produced the rhesus IgG1 recombinant anti-CD8 alpha [MT807R1] mAb (NNHPRR Cat#PR-0817, RRID:AB_2716320), the rhesus IgG1 recombinant anti-CD8 beta [CD8b255R1] mAb (NNHPRR Cat# PR-2557, RRID:AB_2716321), the anti-CD4 [CD4R1] mAb (NNHPRR Cat#PR-0407, RRID:AB_2716322), and the recombinant anti-CD3 [C207]-Diphtheria Toxin mAb (NNHPRR Cat#PR-0307, RRID:AB_2819336). Animals were sedated as per facility standard operating procedures in conjunction with approved VRC animal study protocol. Injection site was shaved and disinfected with alcohol swab. The IV was placed in the saphenous vein and attached to an infusion pump. Total volume of the mAbs was given at a rate of 2mL per minute. We administered two doses (50 mg/kg each), two weeks apart, of anti-CD8α (clone MT807R1; lot: LH18-13), anti-CD8β (clone CD8β255R1; lot: LH18-08), or anti-CD4 (clone CD4R1; lot: MP19-05) to unvaccinated (n=3/group) and IV BCG-vaccinated (n=3/group) macaques. We also treated two groups of animals, each containing one vaccinated and one unvaccinated animal, with up to two doses of anti-CD3 diphtheria toxin-conjugated antibody (clone C207). Animals that received C207 were split into two treatment courses: one group received a single treatment course consisting of 4 total doses at 50 μg/kg/dose once a day for 4 days and were sacrificed 2 weeks after their first dose and a second group received two treatment courses, where the first treatment course mirrored the first group and the second treatment course, administered two weeks later, consisted of an additional 3 doses at 50 μg/kg/dose once a day for 3 days followed by sacrifice 2 weeks later. Our goal was to mimic the timing of our other depletion studies as closely as possible with the amount of C207 available at the time. Regardless of mAb administered, vaccinated macaques received their first dose of depleting mAb between 14- and 18-weeks post vaccination.

### Multiparameter flow cytometry

Generally, samples were analyzed for antigen-specific T cell responses or cellular composition as they were processed. Notable exceptions were peripheral and lung LNs at necropsy and pre-vaccination PBMCs, which were batch-analyzed from cryopreserved samples. Cryopreserved samples were washed, thawed, and rested overnight in R10 before stimulation (29). For T cell stimulation assays, 1–5 million viable cells were plated in 96-well V-bottom plates (Corning) in R10 and incubated with R10 alone (background) or 20 μg ml^−1^ H37Rv Mtb WCL (BEI Resources) for 2 hours before adding 1X protein transport inhibitor cocktail (eBiosciences). The concentration of WCL was optimized to detect CD4 T cell responses; however, protein antigen stimulation may underestimate CD8 T cell responses. For logistical reasons, cells were stimulated overnight (∼14 hours total) before intracellular cytokine staining. For cellular composition determination, cells were stained immediately ex vivo after processing or after thawing. The following mAb conjugates were used: CD3-FITC (clone FN-18), CD3-APC/Cy7 (clone SP34-2), CD4-BV785 (clone OKT4), CD4-BV785 (clone SK3), CD4-BUV395 (clone L200), CD4-BV785 (clone MT477), CD8α-FITC (clone DK25), CD8α-BUV805 (clone RPA-T8), CD8α-BUV805 (clone SK1), CD8β-BUV805 (clone 2ST.5H7), CCR7-BB700 (clone 3D12), CD45-BV510 (clone D058-1283), CD161-BV605 (clone HP-3G10), CD16-BUV496 (clone 3G8), NKG2A-APC (clone Z199), Vg9-Ax680 (clone 7A5), pan γδ TCR-PE (clone 5A6.E9), CD69-ECD (clone TP1.55.3), CD28-PE/Cy5 (clone CD28.2), HLA-DR-PE/Cy5.5 (clone TU36), CD45RA-PE/Cy7 (clone L48), IFNγ-APC (clone B27), TNF-BV650 (clone Mab11), and IL-2-BV750 (clone MQ1-17H12). MR1 monomer was provided by the NIH Tetramer Core Facility and tetramerized with BV421 in-house. All mAb conjugates, tetramers, and viability dye used were titrated to select the dilution with the best separation between positive and negative events. Generally, cells were stained as follows (not all steps apply to all panels, all are at room temperature): Washed twice with PBS/BSA (0.1%); 30-minute incubation with rhesus MR1 tetramer (NIH Tetramer Core Facility) in PBS/BSA; washed twice with PBS; live/dead stain in PBS for 20 minutes; washed twice with PBS/BSA; incubation with surface marker antibody cocktail in PBS/BSA containing 1× Brilliant Stain Buffer Plus (BD Biosciences) for 20 minutes; washed three times with PBS/BSA (0.1%); 20 minute incubation BD Cytofix/Cytoperm Solution (BD Biosciences); washed twice with Perm/Wash Buffer (BD Biosciences); 30 minute incubation with intracellular antibody cocktail in Perm/Wash Buffer containing 1× Brilliant Stain Buffer Plus; washed three times with Perm/Wash Buffer; resuspension in PBS/PFA (0.05%). Data were acquired on a modified BD FACSymphony and analyzed using FlowJo software (v.9.9.6 BD Biosciences). Gating strategies can be found in Figure 1. All cytokine data presented graphically are background subtracted.

**Figure 1.**
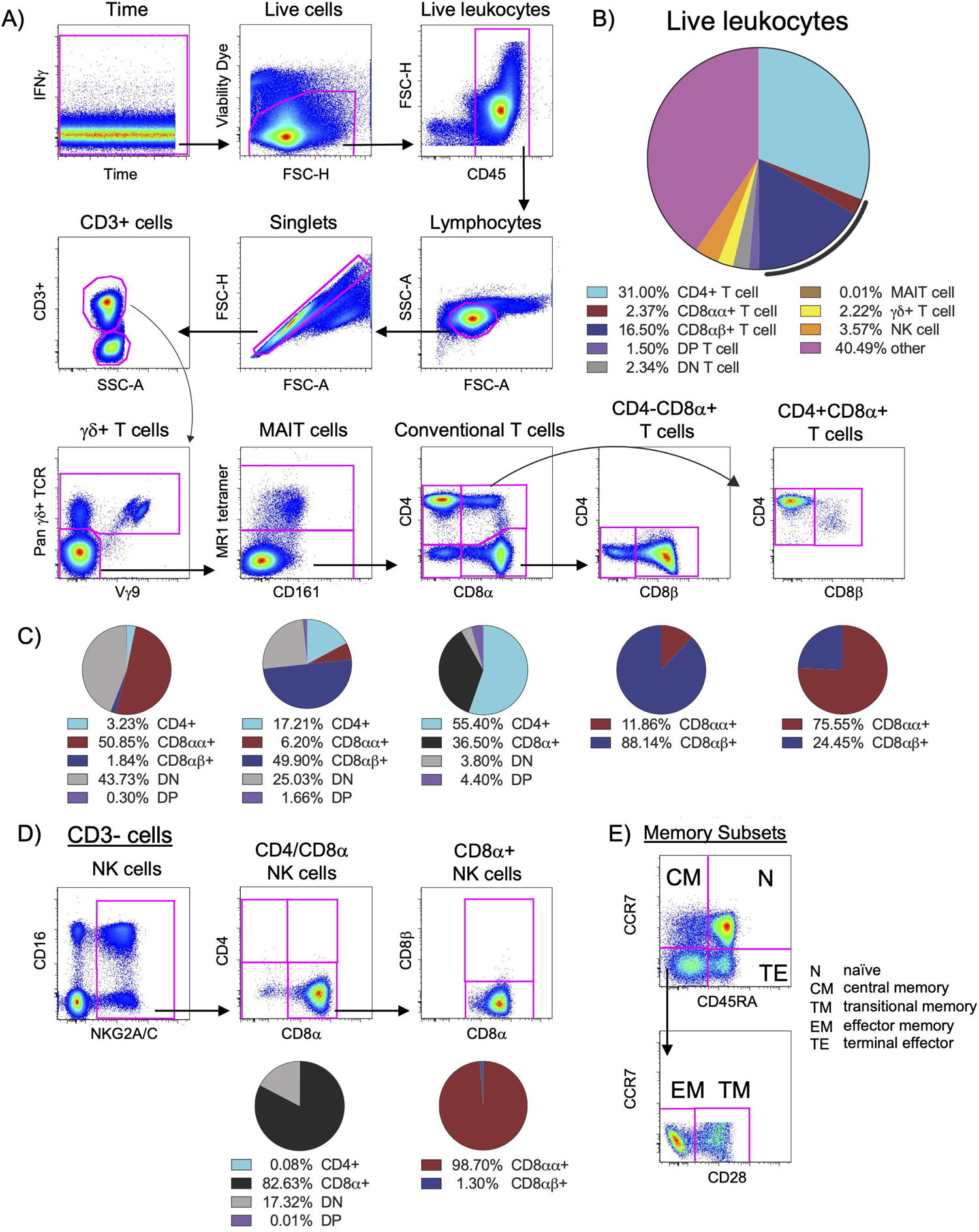
Identification and expression patterns of cell types of interest. (A) Multi-color flow cytometry gating scheme used for identifying CD3+ cell types of interest. (B) Frequencies of cell types of interest as a proportion of live leukocytes. The black line represents total CD8α+ T cells (both CD8αα+ and CD8αβ+). Cells contained within the “other” slice may consist of B cells, monocytes, macrophages, dendritic cells, or stem cells. (C) Expression patterns of CD4, CD8α, and CD8β for γδ+ T cells, MAIT cells, and conventional T cells, as noted. (D) Multi-color flow cytometry gating scheme used for identifying CD3-cell types of interest. The vast majority of NK cells (defined as CD3-NKG2A/C+CD16+, as shown here) express CD8α without co-expression of CD4 nor CD3; thus, we used CD3-CD4-CD8α+ as a surrogate for NK cell quantification. (E) Multi-color flow cytometry gating scheme for identifying memory subsets of conventional T cells. Memory subsets were defined as follows: naïve as CCR7+CD45RA+, central memory as CCR7+CD45RA-, terminal effector as CCR7-CD45RA+, effector memory as CCR7-CD45RA-CD28-, and transitional memory as CCR7-CD45RA-CD28+. Pie charts represent CD4 and CD8 expression patterns from naive PBMC for all 22 animals. Of note, CD8αα and CD8αβ expression was only assessed in animals treated with CD8β255R1.

## Results

### *In vivo* depleting antibodies affect a wide range of lymphocytes

As shown in Figure 1, we targeted for depletion a range of cell types that could potentially have a role in protection, either in IV BCG-vaccinated or unvaccinated animals. We compared the levels of lymphocyte populations following immunodepletion with two doses of anti-CD8α (clone MT807R1; Groups 1, 2), anti-CD8β (clone CD8β255R1, Groups 3, 4), or anti-CD4 (clone CD4R1; Groups 5, 6), given two weeks apart (Figure S1A). Other animals were treated with one to two doses of anti-CD3 diphtheria toxin-conjugated antibody (clone C207 (Groups 7, 8) (Figure S1A). We assessed the longitudinal depletion of lymphocyte subsets, including antigen-specific cells in vaccinated animals, by flow cytometry in PBMCs, BAL, BM, and axillary LNs. We also assessed depletion of individual memory subsets within conventional T cell populations (Figure 1E). At necropsy, we assessed additional peripheral LNs (inguinal and mesenteric), lung-associated LNs (hilar and carinal), lung tissue, and spleen. To properly identify cell types in the presence of saturating levels of depleting mAb, we tested one or more CD4, CD8α, CD8β, or CD3 clones in *ex vivo* staining experiments to identify non-cross-reactive clones that could resolve all cell types (Figure S2).

To interpret *in vivo* cell depletion studies, it is important to consider which cell types express each antigen targeted for depletion. It is also important to consider the relative abundance of individual cell populations as a frequency of all leukocytes (Figure 1B). We first assessed expression patterns of CD4 and CD8α on the surface of conventional CD3+ T cells in PBMCs from unvaccinated macaques. Approximately half of CD3+ T cells expressed only CD4 (light blue) while one third expressed CD8α (black) (Figure 1C). There were also less frequent populations of CD3+ T cells of that do not express CD4 or CD8α (double negative, or DN), or express both CD4 and CD8α (double positive, or DP). Given that surface expression of CD8 can occur as a CD8αα homodimer (red) and CD8αβ heterodimer (dark blue), we evaluated the relative proportions of both types within CD8α+ T cells and within DP T cells (purple). We found the CD8αβ heterodimer most abundant on CD8α+(CD4-) T cells and CD8αα homodimer on DP T cells (Figure 1C). In BAL, approximately half of CD3+ T cells expressed CD8α and one third expressed CD4 (Figure S1B, left). As in PBMCs, the CD8αβ heterodimer in BAL was most abundant on CD8α+(CD4-) T cells (Figure S1B, middle) and DP T cells were predominantly CD8αα+ (Figure S1B, right).

We also assessed expression patterns of CD4, CD8α, and CD8β on the surface of donor unrestricted T cells, such as mucosal associated invariant T (MAIT) cells and γδ+ T cells, as well as natural killer (NK) cells in PBMC (Figure 1C-D) and BAL (Figure S1C) of naïve macaques. In PBMC, nearly half of all MAIT cells expressed CD8β, while CD8β expression was detected on very few γδ+ T cells or NK cells. In contrast, most γδ+ T cells and NK cells expressed CD8αα. Of the three cell types, expression of CD4 was most prevalent on MAIT cells. Co-expression of CD4 and CD8α (DP) was detected rarely on all three cell types; instead, there were sizable proportions of MAIT cells, γδ+ T cells, and NK cells that lacked expression of both CD4 and CD8α (DN) suggesting that sizeable fractions of these cells may not be susceptible to *in vivo* depletion using CD4 or CD8-targeting antibodies. In BAL, we also detected expression of CD8β on a majority of MAIT cells (Figure S1C), though we found an increased prevalence of CD8β on γδ+ T cells and NK cells when compared to PBMC. A majority of γδ+ T cells and NK cells expressed CD8αα in BAL, with a smaller, though sizeable, percentage of MAIT cells expressing CD8αα. Unlike in PBMC, for BAL, we detected populations of MAIT cells that co-expressed both CD4 and CD8α. As in PBMCs, a considerable percentage of γδ+ T cells and NK cells in BAL still lacked expression of CD4 and CD8α, leaving only a very small amount co-expressing both. Thus, in BAL it appears that administration of MT807R1 or CD8β255R1 has the potential to substantially impact levels of MAIT cells and NK cells, though depletion of γδ+ T cells may be more difficult to achieve as they are largely DN T cells.

### Depletion of cell types following MT807R1 treatment

We administered two doses (50 mg/kg each), two weeks apart, of the anti-CD8α depleting antibody, MT807R1, to unvaccinated (n=3) and IV BCG-vaccinated (n=3) macaques and monitored lymphocytes in blood and BAL 2 weeks after each infusion. Here, we report the data in three ways: as frequency of live leukocytes (line graphs), as absolute number after each infusion (line graphs), or as the percent depletion after both infusions (dot plots). Changes to cell population frequencies (LNs and BM) and numbers (PBMCs and BAL) are reported as the median value of all animals between baseline and after two administrations of depleting antibody. Notably, while reductions in many populations were dramatic after the first infusion, the second antibody infusion the administration of a second dose of depleting antibody did not always result in sustained or improved depletion. In PBMCs (Figure 2A), the frequency and absolute cell count were reduced by 1-3 log_10_ for conventional CD8+ T cells (median depletion: 99.5%), DP T cells (median: 95%), MAIT cells (median: 92%), and NK cells (median: 97%). Changes to absolute counts of CD4+ T cells were more variable, with minor to moderate depletion or even increases depending on the animal. Increases in cell number could be due to IL-15-driven homeostatic expansion due to the CD8α depletion (30). Depletion of γδ+ T cells was less complete (median: 53%), most likely because ∼50% of cells do not express CD8α.

**Figure 2.**
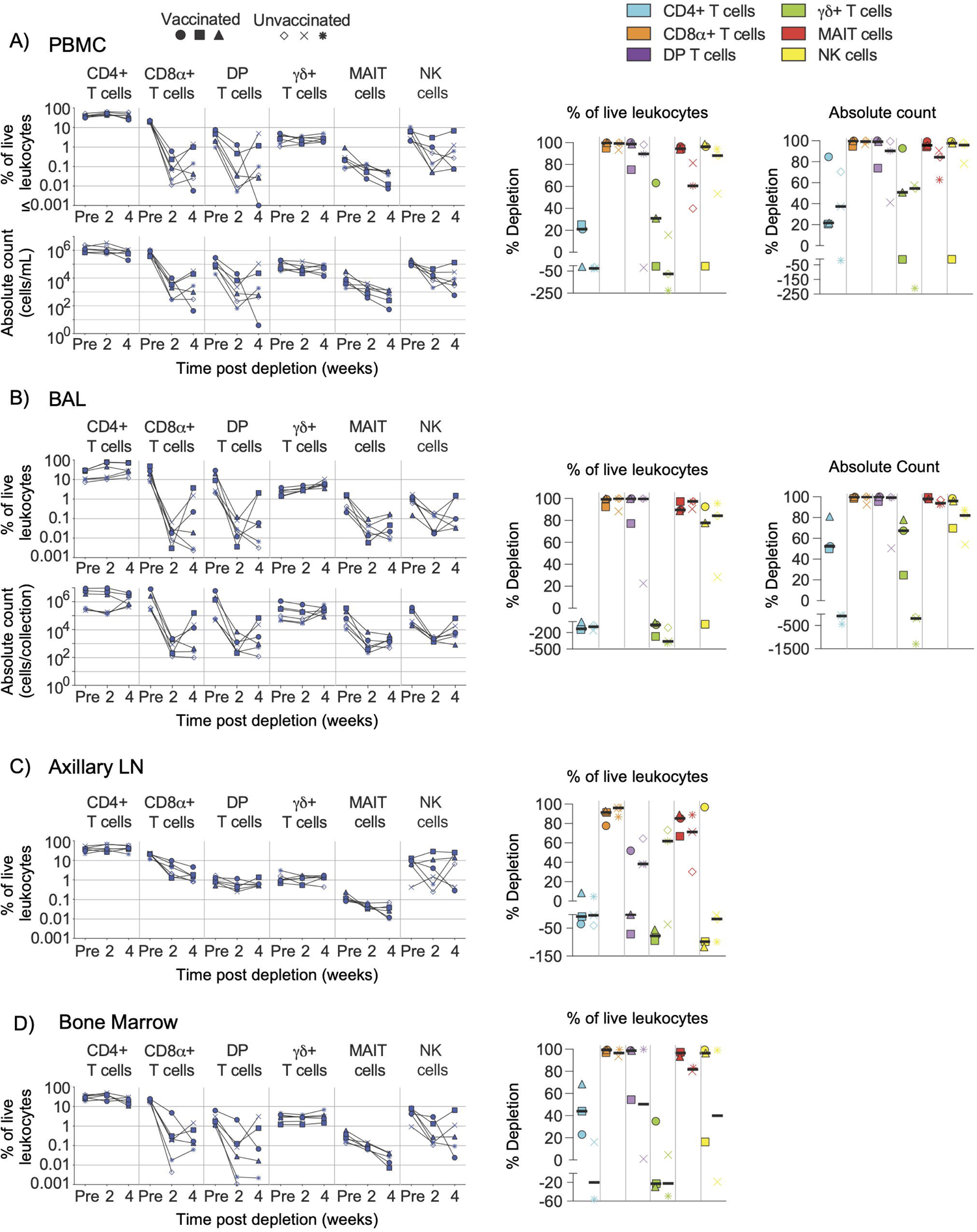
Depletion of select cell types following MT807R1 treatment. Following each administration of MT807R1 we determined the frequency of CD4+ T cells, CD8α+ T cells, DP T cells (CD4+CD8α+), γδ+ T cells, MAIT cells, and NK cells in (A) PBMCs, (B) BAL, (C) axillary lymph node, and (D) bone marrow of vaccinated and unvaccinated macaques. Frequencies were evaluated as a percentage of live leukocytes. We also determined changes in absolute cell counts for each cell type in PBMCs and BAL. We calculated percent depletion before and after two administrations of depleting antibody in two ways: as a proportion of leukocytes and where applicable, using absolute count numbers. In longitudinal graphs and percent depletion dot plots vaccinated animals are displayed as filled symbols and unvaccinated animals as unfilled symbols. Percent depletion plots display each cell type as a unique color according to the legend and a black horizontal line indicates the median value. Negative values in percent depletion plots represent percent increase.

In BAL (Figure 2B), we observed a similar trend in frequency and absolute cell count of leukocytes. Depletions of conventional CD8α+ T cells (median depletion: 99.9%), DP T cells (median: 99.5%), MAIT cells (median: 97%), and NK cells (median: 85%) were highly effective. While the percentage of CD4+ T cells appeared to increase in BAL of all animals, we observed differences in changes to cell count based on vaccination status; vaccinated animals exhibited moderate depletion (median: 52%) while unvaccinated animals exhibited an increase in cell number (median: 91%). Compared to PBMCs, depletion of γδ+ T cells in BAL was more complete, but again we observed a dichotomy based on vaccination status where depletion was observed in vaccinated animals and increases observed in unvaccinated animals.

We also evaluated depletion longitudinally in axillary LNs and BM as a percentage of baseline. In LNs (Figure 2C), depletion of conventional CD8+ T cells (median: 92%) was more pronounced than of DP T cells (median: 38%), while depletion of MAIT cells was intermediate (median: 78%). CD4+ T cells remained unchanged and γδ+ T cells and NK cells increased by ∼50% despite sizeable fractions expressing CD8α. In BM (Figure 2D), depletion was substantial for conventional CD8+ T cells, DP T cells, MAIT cells and NK cells. Minor depletion of CD4+ T cells was also observed and, like other tissue compartments, depletion of γδ+ T cells was lacking; instead, we observed a minor increase.

### Depletion of cell types following CD8β255R1 treatment

An alternate approach for achieving *in vivo* depletion of CD8-expressing cells is the administration of the anti-CD8β depleting antibody, CD8β255R1. Unlike the anti-CD8α depleting antibody MT807R1 that targets cells expressing either CD8αα homodimer or CD8αβ heterodimer, CD8β255R1 depletes only the latter. As such, CD8β255R1 treatment is expected to preferentially deplete conventional CD8+ T cells—the predominant subset that expresses mostly CD8αβ. However, our assessment of CD8α and CD8β expression on the far less predominant donor-unrestricted T cell subsets (Figure 1) suggests that a portion of these cells may also be affected. We assessed depletion in various tissues over 4 weeks following treatment. Changes to cell population numbers (PBMCs and BAL) and frequencies (LNs and BM) are reported as the median value of all animals between baseline and after two administrations of depleting antibody (Figure 3). Of note, for these depletion studies we also differentiated surface expression of the CD8αα homodimer and CD8αβ heterodimer to evaluate the specificity of CD8β depletion (Figure S3). In PBMC, treatment with CD8β255R1 had the greatest effect on conventional CD8+ T cells (median depletion: 93%) and MAIT cells (median: 84%) (Figure 3A). As expected, depletion was specific to only the CD8αβ+ subset of both cell types (median: 99.8%), while the CD8αα+ subset of both cell types remained relatively unchanged (CD8αα+ T cells median: 18%; MAIT cells median: 47%) (Figure S3A). Total DP T cells and γδ+ T cells were only moderately depleted, though substantial reductions were observed in the CD8αβ+ subset (DP T cells median: 99%; γδ+ T cells median: 89%). We observed minimal depletion of CD4+ T cells (median: 35%) and NK cells (median: 50%) following treatment with CD8β255R1.

**Figure 3.**
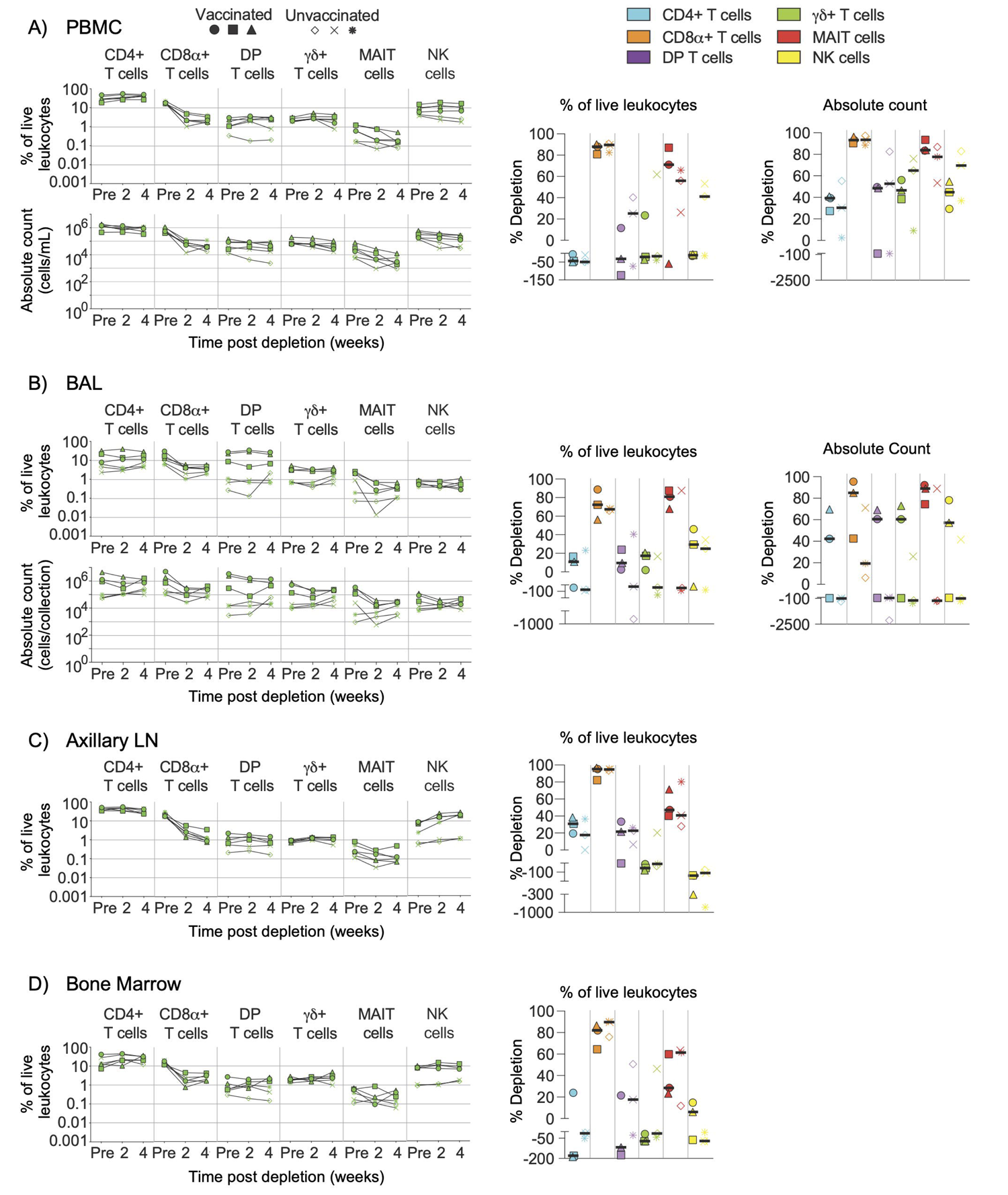
Depletion of select cell types following CD8β255R1 treatment. Following each administration of CD8β255R1 we determined the frequency of CD4+ T cells, CD8α+ T cells, DP T cells (CD4+CD8α+), γδ+ T cells, MAIT cells, and NK cells in (A) PBMCs, (B) BAL, (C) axillary lymph node, and (D) bone marrow of vaccinated and unvaccinated macaques. Frequencies were evaluated as a percentage of live leukocytes. We also determined changes in absolute cell counts for each cell type in PBMCs and BAL. We calculated percent depletion before and after two administrations of depleting antibody in two ways: as a proportion of leukocytes and where applicable, using absolute count numbers. In longitudinal graphs and percent depletion dot plots vaccinated animals are displayed as filled symbols and unvaccinated animals as unfilled symbols. Percent depletion plots display each cell type as a unique color according to the legend and a black horizontal line indicates the median value. Negative values in percent depletion plots represent percent increase.

In BAL, we observed only moderate depletion of total CD8α+ T cells (median: 57%; Figure 3B) while the CD8β+ subset was considerably depleted (median: 97%; Figure S3B). We also observed noticeable increases in the CD8αα+ subset, particularly in unvaccinated animals (Figure S3B). Depletion of DP T cells was confined to the CD8αβ+ subset, while changes to CD8αα+ or CD8αβ+ subsets of γδ+ T cells were highly variable (Figure S3B). Treatment with CD8β255R1 moderately reduced total MAIT cells (median: 82%), though upwards of a 4 log_10_ reduction of the CD8αβ+ subset was observed in some animals. Moderate to minimal increases in cell number of CD4+ T cells and NK cells were observed, respectively (Figure 3B).

In axillary LN (Figure 3C) and BM (Figure 3D), median depletion of total CD8α+ T cells was 95% and 84%, respectively. Again, we observed specific depletion of CD8αβ+ subsets while CD8αα+ subsets remained relatively unchanged (Figures S3C-D). Total DP T cells were only marginally affected by treatment with CD8β255R1. Depletion of DP T cells was most noticeable within the CD8αβ+ subset which only comprises ∼25% of this compartment (Figure 1). The same was true for MAIT cells, where depletion of total MAIT cells was only minimal, with more substantial depletion observed in the CD8αβ+ subset (Figures S3C-D). In both LN and BM, total γδ+ T cells were relatively unaffected by treatment with CD8β255R1, even the CD8αβ+ subset in the BM (Figure S3D), though improved depletion of the CD8αβ+ subset was observed in LN (Figure S3C).

### Depletion of cell types following CD4R1 treatment

We quantified subsets following administration of CD4R1 that targets CD4+ T cells. Changes to cell population frequencies (LNs and BM) and numbers (PBMCs and BAL) are reported as the median value of all animals between baseline and after two administrations of depleting antibody. In PBMCs (Figure 4A), we observed substantial depletion of CD4+ T cells (median: 98%) and moderate depletion of DP T cells (median: 89%). Changes to MAIT cell counts were variable and ranged from 5% to 91% depletion depending on the animal. We observed minimal changes to CD8+ T cells (median: 33% depletion) and γδ+ T cells (median: 38% increase), while a pronounced increase was observed in NK cells (2.5-fold).

**Figure 4.**
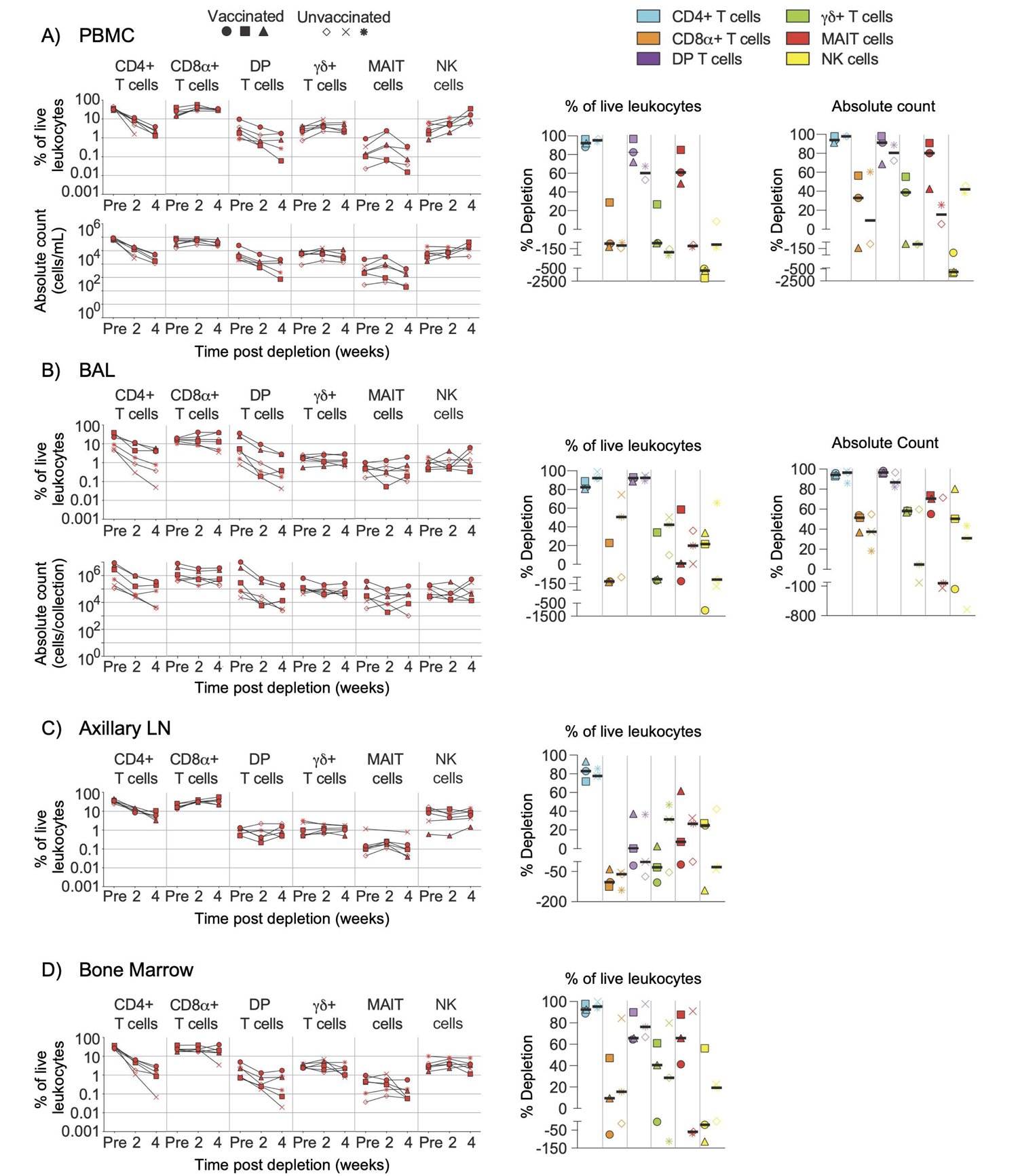
Depletion of select cell types following CD4R1 treatment. Following each administration of CD4R1 we determined the frequency of CD4+ T cells, CD8α+ T cells, DP T cells (CD4+CD8α+), γδ+ T cells, MAIT cells, and NK cells in (A) PBMCs, (B) BAL, (C) axillary lymph node, and (D) bone marrow of vaccinated and unvaccinated macaques. Frequencies were evaluated as a percentage of live leukocytes. We also determined changes in absolute cell counts for each cell type in PBMCs and BAL. We calculated percent depletion before and after two administrations of depleting antibody in two ways: as a proportion of leukocytes and where applicable, using absolute count numbers. In longitudinal graphs and percent depletion dot plots vaccinated animals are displayed as filled symbols and unvaccinated animals as unfilled symbols. Percent depletion plots display each cell type as a unique color according to the legend and a black horizontal line indicates the median value. Negative values in percent depletion plots represent percent increase.

In BAL (Figure 4B), depletion of CD4+ T cells (median: 95%) was comparable to that observed in PBMCs, while DP T cells exhibited better depletion (median: 96%). Changes in MAIT cell count differed by vaccination status, with moderate depletion occurring in all vaccinated animals (median: 71%), while unvaccinated animals exhibited a range of outcomes from 71% depletion to 140% increase. Increases of CD8+ T cells, γδ+ T cells, or NK cells was not observed in BAL as it was in PBMCs; instead, only minor to moderate depletion was observed in most animals.

LN biopsies revealed a moderate depletion of CD4+ T cells (Figure 4C), though two administrations of CD4R1 did not appear to alter the frequency of DP T cells. CD4R1 did not appear to substantially alter γδ+ T cells, MAIT cells, or NK cells. Notably, both vaccinated and unvaccinated animals exhibited a moderate increase to CD8+ T cell frequency. Compared to the LN, the depletion that followed administration of two CD4R1 doses was more complete in BM (Figure 4D). Indeed, a 1-2 log_10_ reduction was observed in CD4+ T cells, though reductions in the frequency of DP T cells and γδ+ T cells was less complete. We observed a high degree of variability of MAIT cell frequencies based on vaccination status. All vaccinated animals exhibited moderate depletion of MAIT cells while unvaccinated animals ranged from 91% depletion to 68% increase. Changes in NK cell frequencies were similarly variable in vaccinated animals and ranged from 56% depletion to 110% increase. Unvaccinated animals exhibited minimal alteration of NK cell frequencies.

### Depletion of select cell types following DT390-scfbDb (C207) treatment

Despite numerous studies using mAbs for selective *in vivo* cell depletion, the precise mechanisms underlying the depletion remains unclear; also, most mAbs against cell surface antigens do not deplete cells for reasons that are ill-understood. An alternative approach to achieve *in vivo* depletion is to use mAbs, or their fragments, that have been chemically cross-linked with an immunotoxin, such as diphtheria toxin. One such molecule, anti-CD3 [C207]-diphtheria toxin, has shown enhanced bioactivity when in a fold-back single-chain diabody format and has successfully been used in NHP models (11–13). To assess *in vivo* depletion by C207, we infused it daily into NHPs, giving either one or two courses of treatment depending on animal group (Figure S1A).

In PBMCs (Figure 5A) and BAL (Figure 5B), we observed considerably less depletion of CD4, CD8α+, and DP T cells when compared to MT807R1, CD8β255R1, and CD4R1. In contrast, treatment with C207 resulted in substantial depletion of MAIT cells in PBMCs that was comparable to animals that received MT807R1. This was not the case in BAL, where treatment with C207 resulted in only moderate depletion. Notably, we observed the most complete depletion of γδ+ T cells in PBMCs and BAL following treatment with C207 compared MT807R1, CD8β255R1, and CD4R1.

**Figure 5.**
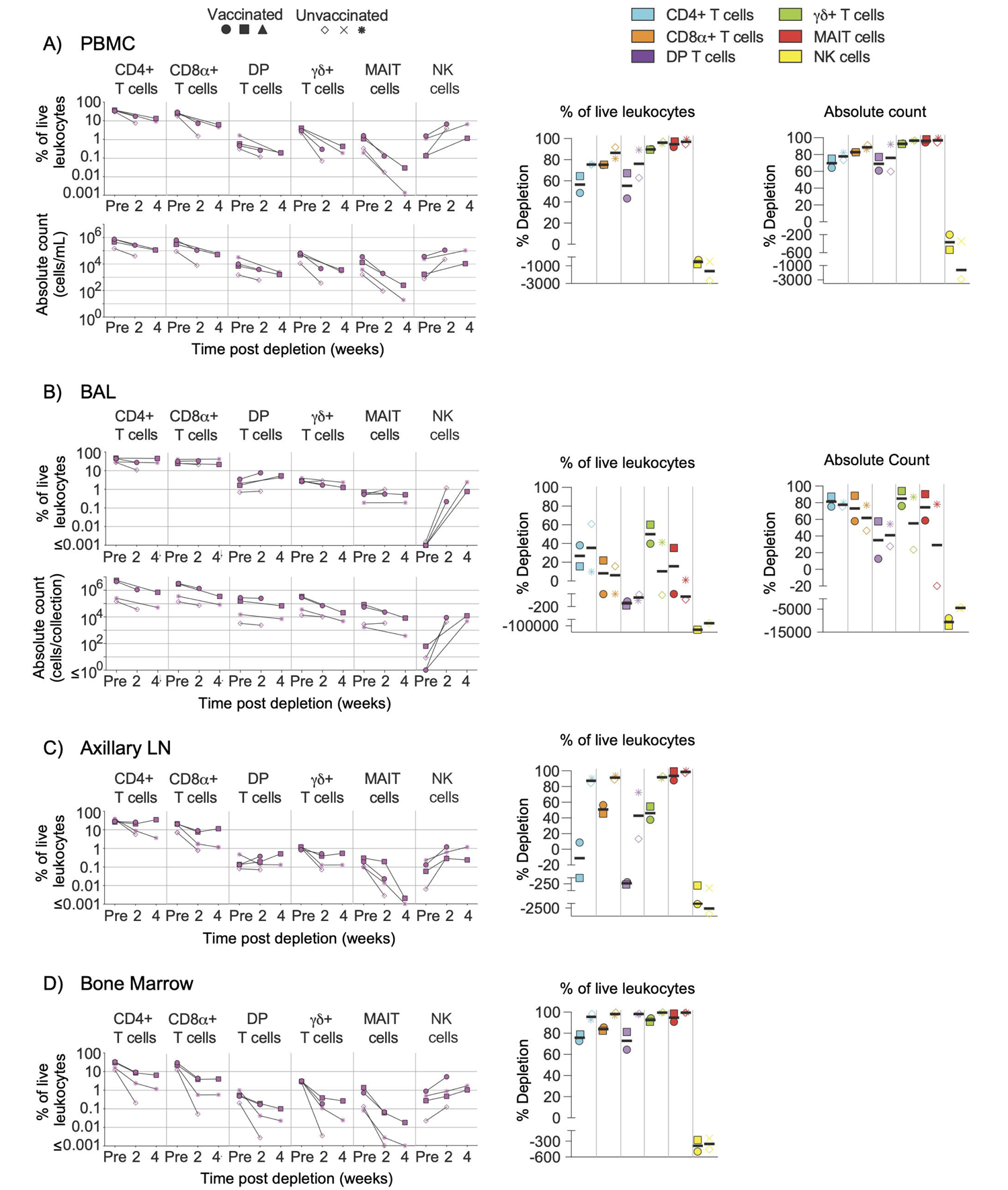
Depletion of select cell types following C207 treatment. Following each administration of C207 we determined the frequency of CD4+ T cells, CD8α+ T cells, DP T cells (CD4+CD8α+), γδ+ T cells, MAIT cells, and NK cells in (A) PBMCs, (B) BAL, (C) axillary lymph node, and (D) bone marrow of vaccinated and unvaccinated macaques. Frequencies were evaluated as a percentage of live leukocytes. We also determined changes in absolute cell counts for each cell type in PBMCs and BAL. We calculated percent depletion before and after two administrations of depleting antibody in two ways: as a proportion of leukocytes and where applicable, using absolute count numbers. In longitudinal graphs and percent depletion dot plots vaccinated animals are displayed as filled symbols and unvaccinated animals as unfilled symbols. Percent depletion plots display each cell type as a unique color according to the legend and a black horizontal line indicates the median value. Negative values in percent depletion plots represent percent increase. Of note, one pair of vaccinated and unvaccinated animals received one dose, while another pair received up to 2 doses.

LN biopsies revealed few changes to the frequency of conventional T cells following treatment with C207 (Figure 5C). Of note, this was exclusive to vaccinated animals while unvaccinated animals achieved up to 1 log_10_ depletion for the same cell types. The same observation was made in γδ+ T cells. Notably, treatment with C207 substantially depleted MAIT cells, while also resulting in a large increase in the frequency of NK cells. Analysis of BM aspirates revealed up to a 3 log_10_ reduction in the frequency conventional T cells, as well as γδ+ T cells and MAIT cells, both in vaccinated and unvaccinated animals (Figure 5D). In agreement with our other findings, the frequency of NK cells increased considerably in the BM following C207 treatment.

### Depletion of memory subsets and antigen-specific cell populations

Depletion of differentiation-defined subsets of conventional T cells (CD8+, CD4+, and DP) in PBMCs and BAL were also evaluated in all animals (Figure S4), as were antigen-specific cells in vaccinated macaques (Figure 6). For memory T cell subsets, we evaluated naïve, central memory, transitional memory, effector memory, and terminal effector T cells using combinations of CCR7, CD45RA, and CD28 expression (Figure 1). In PBMCs and BAL, differentiation status did not affect the extent to which depletion of targeted cell types occurred. Some minor exceptions to this finding were less effective depletion of effector memory DP T cells in PBMCs of unvaccinated animals that received MT807R1 (Figure S4A), and less effective depletion of transitional memory CD4+ T cells in vaccinated animals that received CD4R1 (Figure S4C). In animals that received CD8β255R1, depletion of CD8+ T cell memory subsets in PBMC and BAL was specific to CD8+ T cells and DP T cells that expressed CD8β (Figure S4B). DP T cells in BAL that expressed CD8β also had slight variation in depletion across memory subsets, particularly within the naïve subset of unvaccinated animals. Treatment with C207 also led to similar depletion across most memory subsets (Figure S4D), though we noted some exceptions: in PBMCs of both vaccinated and unvaccinated animals we observed an increase of transitional memory CD4+ T cells, and higher levels of depletion within naïve and terminal effector subsets of both CD8+ T cells in PBMCs and DP T cells in BAL.

**Figure 6.**
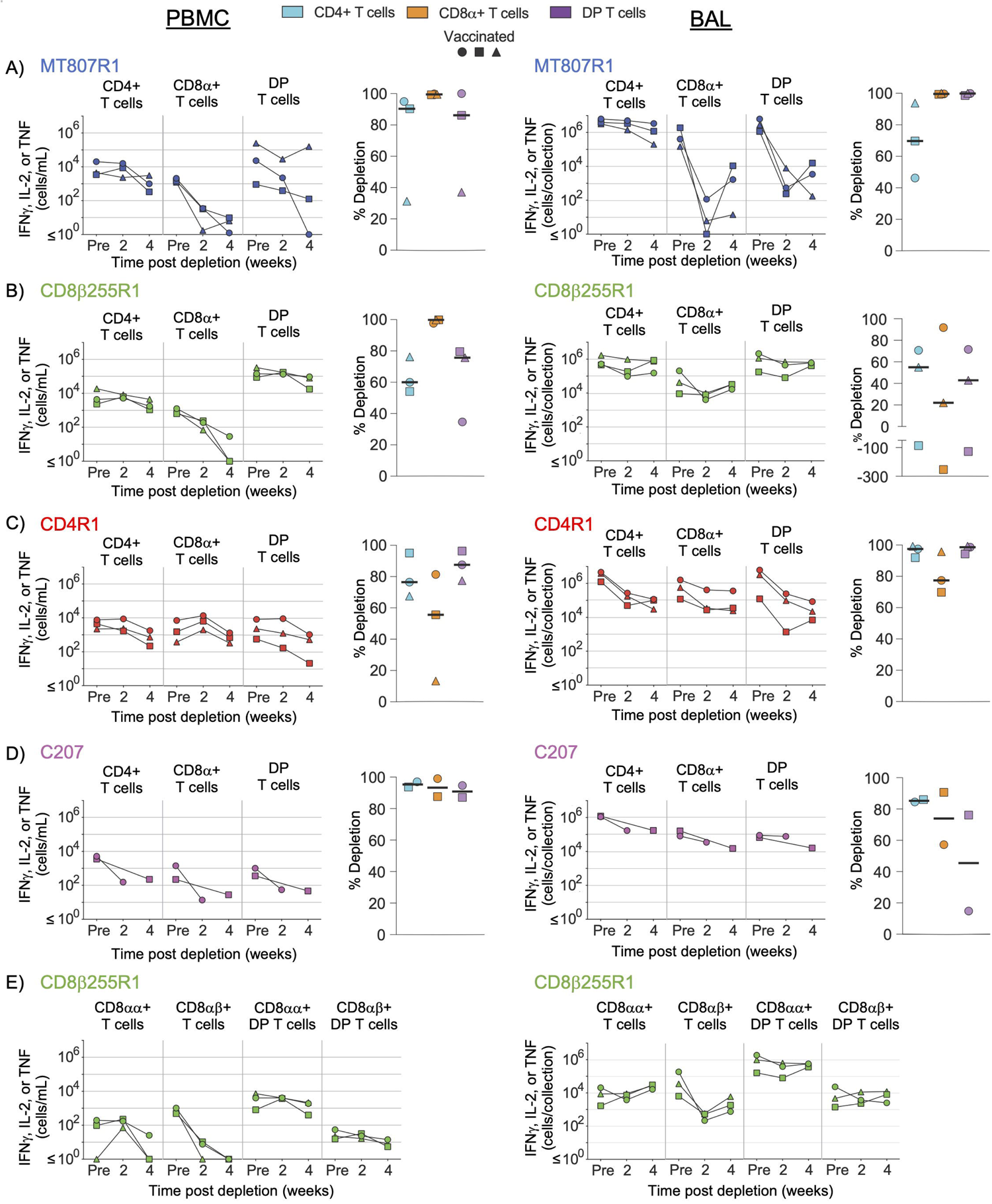
Antigen-specific cells in vaccinated macaques are depleted equally *in vivo*. Depletion of antigen-specific cells, measured by secretion of IFNγ, IL-2, or TNF after stimulation with *Mtb* whole-cell lysate, was assessed in PBMCs (left) and BAL (right) of vaccinated macaques that received (A) MT807R1, (B) CD8β255R1, (C) CD4R1, or (D) C207. In animals that received CD8β255R1, we also evaluated antigen-specific CD8+ and DP T cells (CD4+CD8α+) that specifically expressed CD8β (E). Vaccinated animals are displayed as filled symbols. Percent depletion plots display each cell type as a unique color according to the legend and a black horizontal line indicates the median value. Negative values in percent depletion plots represent percent increase.

In vaccinated NHPs, we also evaluated depletion of antigen-specific CD4+, CD8+, and DP T cells in PBMCs and BAL, defined as cytokine-expressing T cells, following restimulation with *Mtb* whole-cell lysate (Figure 6). In animals that received MT807R1, depletion of antigen-specific CD8+ T cells was nearly complete, while antigen-specific DP T cells and CD4+ T cells were moderately depleted in two out of three animals (Figure 6A, left). Similar observations were made in BAL, though depletion of antigen-specific CD4+ T cells was less substantial (Figure 6A, right). In animals that received CD8β255R1, we observed more complete depletion of antigen-specific CD8+ T cells in PBMCs (Figure 6B, left) than in BAL (Figure 6B, right). A similar observation was made when assessing CD8β-expressing CD8+ and DP T cells in PBMCs and BAL (Figure 6E). Of note, we observed depletion of CD8αα+ T cells in PBMCs following CD8β255R1 treatment, and lack of depletion of CD8β-expressing DP T cells in both PBMCs and BAL in most animals. Antigen-specific CD4+ T cells were depleted variably in PBMCs and BAL of animals that received CD8β255R1.We observed higher levels of depletion for antigen-specific CD4+ T cells in BAL (Figure 6C, right) than in PBMCs (Figure 6C, left) of animals that received CD4R1, while moderate depletion of CD8+ T cells was observed in both PBMCs and BAL. Lastly, depletion of antigen-specific CD4+ T cells, CD8+ T cells, and DP T cells in animals that received C207 was particularly effective in the PBMC although much less so in BAL (Figure 6D). In many cases, the trends observed in antigen-specific cells were similar to that observed in non-antigen specific cells.

### Comparison of antibody-mediated depletion across multiple tissues

The unrestricted nature of CD4, CD8α, and CD8β expression on multiple cell types presents a challenge when evaluating the importance of an individual cell type during *in vivo* depletion studies. Treatment with MT807R1, CD8β255R1, CD4R1, or C207 provided us the unique opportunity to compare across depletion groups the extent by which depletion was observed in multiple tissues at necropsy. For PBMCs (Figure S5A), BAL (Figure S5B), and BM (Figure S5C) we compiled frequency and absolute count data from Figures 2-6 for ease of comparison across treatment groups. Treatment with MT807R1 or CD8β255R1 resulted in comparable depletion of total CD8+ T cells in BAL, though MT807R1 depleted them better in PBMCs and BM. We observed the lowest γδ+ T cell count in PBMCs and BM for animals that received C207, while similar levels were observed in animals that received CD4R1 in BAL.

We also evaluated depletion in additional peripheral LNs (mesenteric and inguinal), lung LNs (hilar and carinal), spleen, and up to 6 lung lobes at necropsy. In lung lobes (Figure 7A), we observed similar frequencies of CD4+ T cells in individual lung lobes of animals that received CD4R1 or C207, though only animals that received C207 exhibited low CD4+ T cell counts. Depletion of total CD8α+ T cells was most pronounced following treatment with MT807R1 and C207. DP T cell counts were lowest in the lung lobes of animals that received C207, though DP T cells were also well depleted in many MT807R1-treated animals and one CD4R1-treated animal. In the lung lobes of animals that received CD8β255R1, the frequency of CD8α+ T cells was higher than in animals that received MT807R1. However, this observation was not indicative of overall cell numbers, as CD8α+ T cell counts in lung lobes of CD8β255R1-treated animals were often comparable to those of CD4R1-treated animals. As expected, the frequency and total count of NK cells was lower in MT807R1-treated animals compared to CD8β255R1-treated animals due to the majority of these cells predominantly expressing CD8αα. Despite a large proportion of MAIT cells in BAL expressing CD8β (Figure 1), treatment with CD8β255R1 did not appreciably deplete them in lung lobes. γδ+ T cells were lowest in C207-treated animals. However, while reductions in frequency of cell types can appear complete, such as with MAIT cells in our MT807R1-treated group, substantial numbers of cells can remain in the tissues.

**Figure 7.**
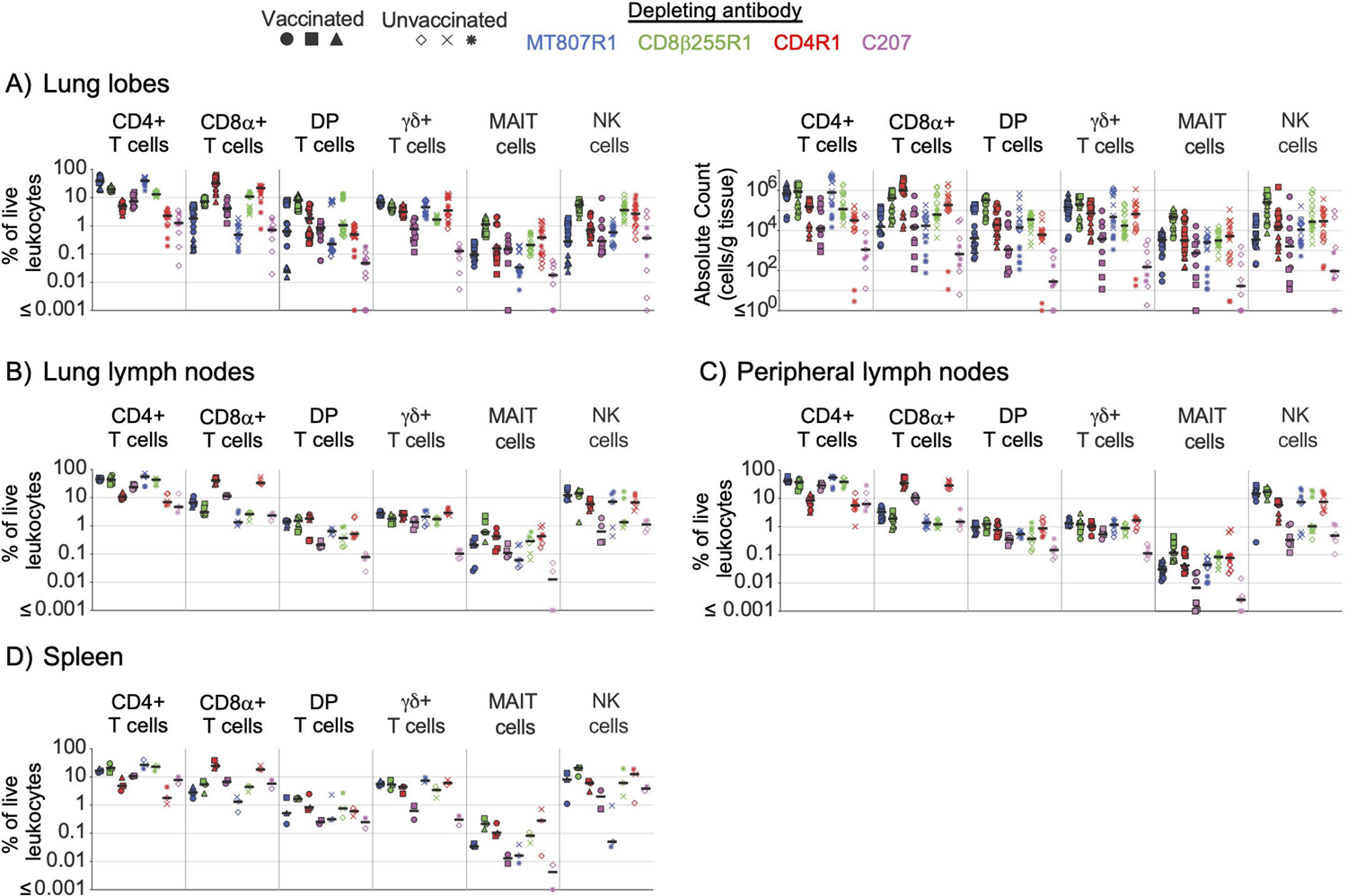
Comparison of antibody-mediated depletion in multiple tissues. Comparison of cell frequency (percentage of live leukocytes) of CD4+ T cells, CD8α+ T cells, DP T cells (CD4+CD8α+), γδ+ T cells, MAIT cells, and NK cells in (A) lung lobes, (B) lung lymph nodes, (C) peripheral lymph nodes, and (D) spleen of vaccinated and unvaccinated animals following antibody-mediated depletion. Lung lymph nodes include hilar and carinal lymph nodes while peripheral lymph nodes include axillary, inguinal, and mesenteric lymph nodes. Lung lobes (up to 6 total) and lymph nodes are shown individually for each animal. Comparisons of absolute cell counts are shown for lung lobes. Vaccinated animals are denoted by filled symbols and unvaccinated animals by unfilled symbols. Animals that received MT807R1 are shown in blue, CD8β255R1 are shown in green, CD4R1 are shown in red, and C207 are shown in pink. Median values are shown with a black horizontal line.

From hilar and carinal LNs within the lung (Figure 7B) we observed the lowest frequency of CD4+ T cells in animals treated with CD4R1 or C207. Animals treated with MT807R1, CD8β255R1, or C207 had similar levels of CD8α+ T cells, all of which were lower compared to those that received CD4R1. In contrast to observations in the lung lobes, frequencies of DP T cells in animals treated with CD8β255R1 were comparable to animals treated with MT807R1 or CD4R1 and lowest in animals that received C207. γδ+ T cell frequencies were also not reduced in C207-treated animals, as in lung lobes, but were instead comparable in all groups. Frequencies of MAIT cells in lung lymph nodes mirrored our observations in lung lobes, though MT807R1-treatment was less effective at reducing frequencies of NK cells in LNs within the lung. We observed similar trends in cell frequencies from axillary, mesenteric, and inguinal LNs within the periphery (Figure 7C), but with a subset of C207-treated animals exhibiting reduced levels of MAIT cells when compared to MT807R1-treated animals. Of note, peripheral LN comparisons include necropsy data from axillary lymph nodes that were presented in Figures 2-5. With few exceptions, the trends we observed in lymph nodes further extended to our observations in BM (Figure S5C) and spleen (Figure 7D). Of note, BM and spleen CD8α+ and DP T cell frequencies were lower in MT807R1-treated animals compared to those that received CD8β255R1, and C207 treatment resulted in the lowest levels of γδ+ T cells. Importantly, in the case where multiple samples were collected from the same anatomical site, such as with lung lobes or multiple lung-associated or peripheral LNs, comparable depletion was frequently observed.

We also compared depletion of antigen-specific cells in multiple tissues across depletion groups at necropsy. Comparisons across groups at necropsy for PBMCs (Figure S6A) and BAL (Figure S6B) were compiled from data presented in Figure 6. Of note, depletion of antigen-specific CD8α+ T cells in PBMCs was most complete in animals that received CD8β255R1, though this was not the case in BAL where the most complete depletion was observed in animals that received MT807R1. In animals that received CD4R1, antigen-specific CD4+ T cell counts were lowest in BAL while cell counts in PBMCs were comparable to animals that received MT807R1 or CD8β255R1. Antigen-specific DP T cell numbers in PBMCs were lowest in animals that received MT807R1 or C207, while numbers in BAL were lowest in animals that received MT807R1.

For lung lobes (Figure 8A), we observed lower numbers of antigen-specific CD8α+ T cells in animals that received MT807R1, CD8β255R1, or C207 when compared to animals that received CD4R1. In some cases, this depletion was essentially complete with less than 1 cell per gram of tissue. In contrast, while antigen-specific CD4+ T cell numbers were lower in animals that received CD4R1 compared to all other groups, the absolute count of these cells still ranged from 10^4^-10^5^ cells per gram of tissue. For antigen-specific DP T cells in the lung, we observed the lowest numbers in animals that received C207, followed by animals that received MT807R1. In lung LNs (Figure 8B), we were surprised to observe a lower frequency of antigen-specific CD4+ T cells in animals that received MT807R1 or CD8β255R1 compared to those that received CD4R1. Further, the frequency of antigen-specific CD8α+ T cells were comparable in all groups except C207, where they were substantially lower. Antigen-specific cells in peripheral LNs (Figure 8C) were considerably lower than what we observed in lung LNs, and followed the trends we observed in lung tissue apart from overall lower frequencies for all cell types. In BM (Figure 8D), animals that received C207 appeared to generally have lower frequencies of antigen-specific CD4+, CD8α+, and DP T cells, though we did observe a few exceptions. Similar to observations in lung LNs, antigen-specific cells in the spleen (Figure 8E) were comparable across animals that received MT807R1, CD8β255R1, and CD4R1. Animals that received C207 had the lowest frequency of antigen-specific CD4+ and DP T cells while CD8α+ T cells varied.

**Figure 8.**
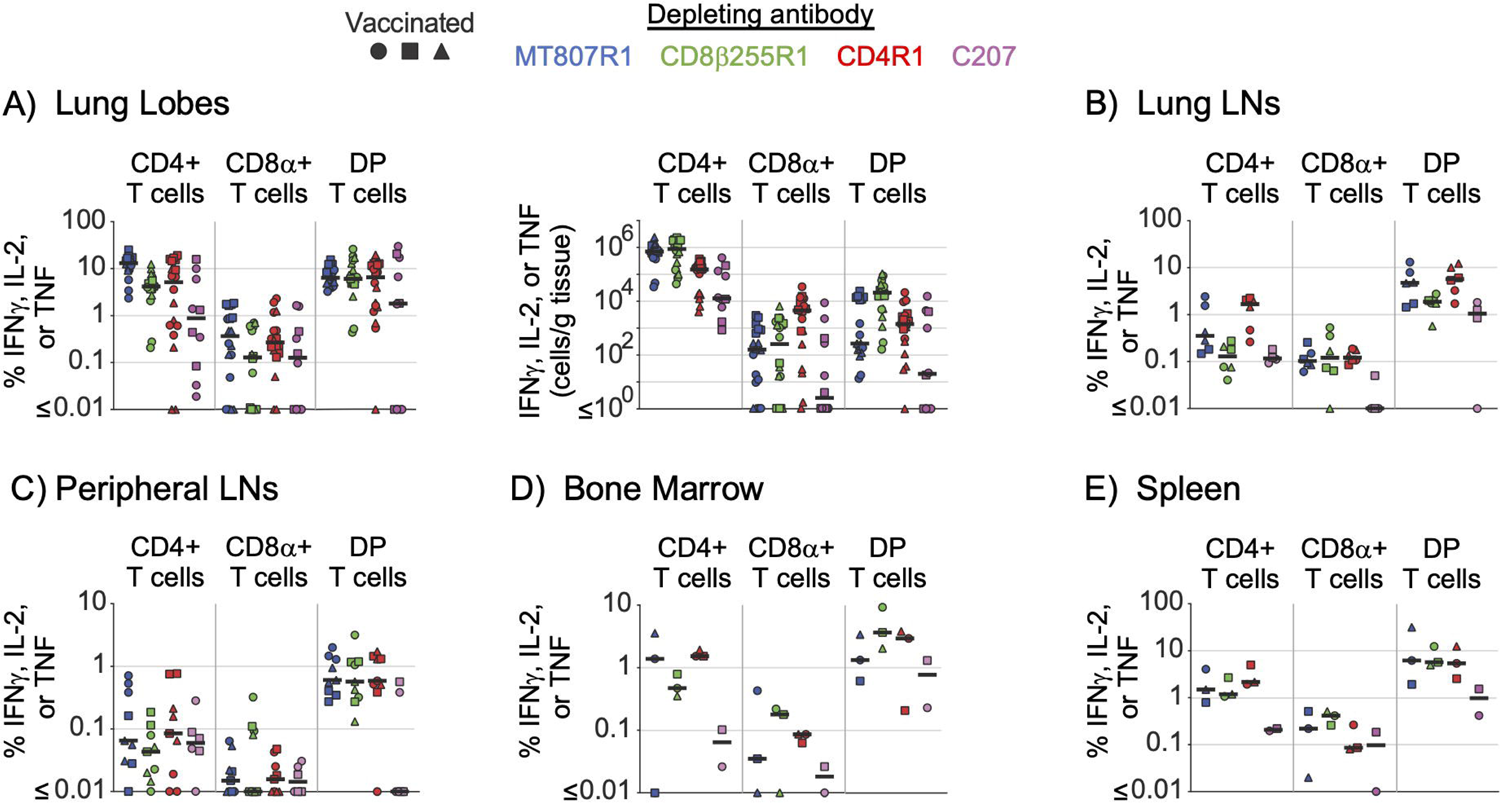
Comparison of antibody-mediated depletion of antigen-specific cells in multiple tissues. Comparison of antigen-specific cell frequency (percentage of CD4+, CD8+, or DP T cells, as noted), measured by secretion of IFNγ, IL-2, or TNF after stimulation with *Mtb* whole-cell lysate, of CD4+ T cells, CD8α+ T cells, and DP T cells (CD4+CD8α+) in (A) lung lobes, (B) lung lymph nodes, (C) peripheral lymph nodes, (D) bone marrow, and (E) spleen of vaccinated animals following antibody-mediated depletion. Lung lymph nodes include hilar and carinal lymph nodes while peripheral lymph nodes include axillary, inguinal, and mesenteric lymph nodes. Lung lobes are shown individually (up to 6 total) for each animal. Comparisons of absolute cell counts are shown for lung lobes. Vaccinated animals are denoted by filled symbols. Animals that received MT807R1 are shown in blue, CD8β255R1 are shown in green, CD4R1 are shown in red, and C207 are shown in pink. Median values are shown with a black horizontal line.

## Discussion

Specificity and affinity for only its target molecule makes mAbs attractive options for *in vivo* depletion studies. However, it is not uncommon for more than one cell type to share expression of the same surface marker that would be unintended targets of a depleting mAb (31). For instance, an anti-CD8α depleting mAb will not only target conventional CD8αβ+ T cells, but also smaller populations of other less prevalent CD8α-expressing cells such as DP T cells, γδ cells, MAIT cells, and some CD3-cells including NK cells. Indeed, we observed up to a 3 log_10_ reduction in absolute count for these CD8α-expressing cell types in PBMCs and BAL of animals that received MT807R1. While an anti-CD8β depleting mAb has increased specificity for conventional CD8+ T cells, since most express the CD8αβ heterodimer, we still observed considerable depletion of CD8αβ-expressing MAIT cells, DP cells, and γδ+ T cells in PBMC and BAL. Furthermore, we observed substantial numbers of conventional CD8+ T cells, expressing only the CD8αα homodimer, that remained in PBMCs and BAL after CD8β antibody infusion. Thus, complete depletion of conventional CD8+ T cells, and only these cells, using either an anti-CD8α or anti-CD8β depleting antibody is not possible with available reagents.

The objective of *in vivo* depletion studies is the substantial or complete removal of a select cell type throughout the entire body. Incomplete depletion may make it difficult to attribute a particular outcome to the absence of that cell type. Whether these residual cells meaningfully contribute to biological function must be evaluated in any given application. It is also important to recognize that depletion studies such as these have profound influence on the homeostatic control of cell populations, often leading to increases in untargeted cell populations.

Antibody-mediated depletion has generally been most successful in peripheral blood and more difficult to achieve in tissue (16, 32, 33). In PBMCs, treatment with MT807R1 or CD8β255R1 resulted in the most extensive depletion of targeted cells, followed by treatment with CD4R1, and finally with C207. Interestingly, despite a high proportion of γδ+ T cells expressing CD8α, they were largely unaffected by treatment with MT807R1. While treatment with CD8β255R1 did result in depletion of CD8β-expressing γδ+ T cells, their removal did not noticeably affect the total γδ+ T cell count, possibly due to inaccuracies in enumerating the small size of this CD8β-expressing subset. Appreciable depletion of total γδ+ T cells was observed only in animals that received immunotoxin C207.

Vaccine-elicited cells – which may be activated, more numerous, or resident in specific tissues– may differ in resistance to antibody-mediated depletion compared to resting, recirculating populations. To address this, we used macaques immunized intravenously with BCG, as these animals have previously been shown to exhibit a large increase in activated T cell numbers in BAL and lung tissue. In BAL of vaccinated macaques, effector memory and terminal effector conventional T cells (CD4+, CD8+, and DP) were depleted comparable to other memory subsets. Depleting mAbs that targeted CD4, CD8α, or CD8β exhibited the most complete depletion, while more variability between memory subsets was observed in animals that received C207. Furthermore, total antigen-specific conventional T cells were best depleted in animals that received MT807R1 or CD4R1. However, we did observe substantial depletion of antigen-specific CD4+ T cells in one animal that received MT807R1. The same observation was made for antigen-specific CD8+ T cells in one animal that received CD4R1. This variability in off-target cell depletion highlights the importance of appropriately powering any depletion studies that aim to understand depletion of antigen-specific T cells as considerable variation in depletion was observed in many subsets.

For all depleting antibodies tested, we consistently observed less depletion in axillary LNs compared to PBMCs. This was particularly noticeable for DP T cells, a small subset of CD3+ T cells that may play a role in the host response to a variety of infections including *Mtb* (*34*). For this subset, depletion reached only 38% in LNs compared to 95% in PBMCs of animals that received MT807R1. This was not the case for all tissues, as we observed substantial depletion of targeted cell types in BAL and BM depending on the depleting mAb given. Furthermore, when we analyzed depletion in individual lung lobes, we generally observed consistent depletion across each, though some variability did exist across animals.

In this study, we extensively evaluated the degree of *in vivo* depletion in multiple tissues of unvaccinated and IV BCG-vaccinated nonhuman primates following two administrations of an anti-CD8α, anti-CD8β, or anti-CD4 depleting mAb. We also explored the impact of up to two administrations of an alternative means of achieving *in vivo* depletion by using an anti-CD3 diphtheria toxin-conjugated mAb. Substantial, though not complete, depletion of targeted cells was observed, however, it varied based on the cell type and anatomical location. Notably, we included both representational (percent of leukocytes), and absolute number, measurements of *in vivo* depletion at various tissue sites to highlight that even when substantial reductions in cell representation occurred, considerable numbers of residual cells remained. Overall, we found that use of an anti-CD8α or anti-CD8β depleting mAb resulted in the most comprehensive depletion of targeted cell types, though neither can remove a single cell type exclusively. Antibody targeting CD4 was almost as good. However, despite the ability to directly mediate cell death via conjugation to diphtheria toxin, the anti-CD3 mAb did not perform as well as any of the three conventional depleting mAbs that were tested, though we did observe improved depletion of γδ+ T cells that were largely unaffected by conventional depleting mAbs. Interpretation of *in vivo* depletion studies warrants careful consideration of target cell selection due to surface marker expression promiscuity, particularly that of CD8α and CD8β, and should be properly powered to account for variability in off-target effects that may occur between animals. Further, variability of the effector mechanism (e.g. FcgR, complement) could also affect depletion and necessitates future studies evaluating these effector mechanisms for individual depleting mAbs. Overall, the use of *in vivo* depleting mAbs is a useful strategy to look at the role of specific cell types though the variability of and extent of depletion can impact interpretation of results.

## Data availability statement

The original contributions presented in the study are included in the article/supplementary material. Further inquiries can be directed to the corresponding author.

## Author Contributions

M.S.S., M.K., C.C.L., S.P., A.N.B., and P.A.D. performed *in vivo* depletion studies. P.A.D., D.M.M., R.S., and M.R. provided experimental oversight. M.S.S. and M.R. wrote the initial draft and generated figures, with the other authors providing comments.

## Acknowledgements

We thank Dr. Elizabeth Lake Potter for data processing assistance, members of the ImmunoTechnology Section for discussion and guidance, the Nonhuman Primate Immunogenicity Core for assistance processing specimens, and the Flow Cytometry Core for expert assistance in instrumentation. The depleting mAbs used in this study were provided by the NIH Nonhuman Primate Reagent Resource (ORIP P40 OD028116, U24 AI126683).

## Supplementary Material

The Supplementary Material for this article can be found online at:

