## Supplementary figures and images for "Antibody-mediated depletion of select T cell subsets in blood and tissue of nonhuman primates"

### Supplemental Figure 1

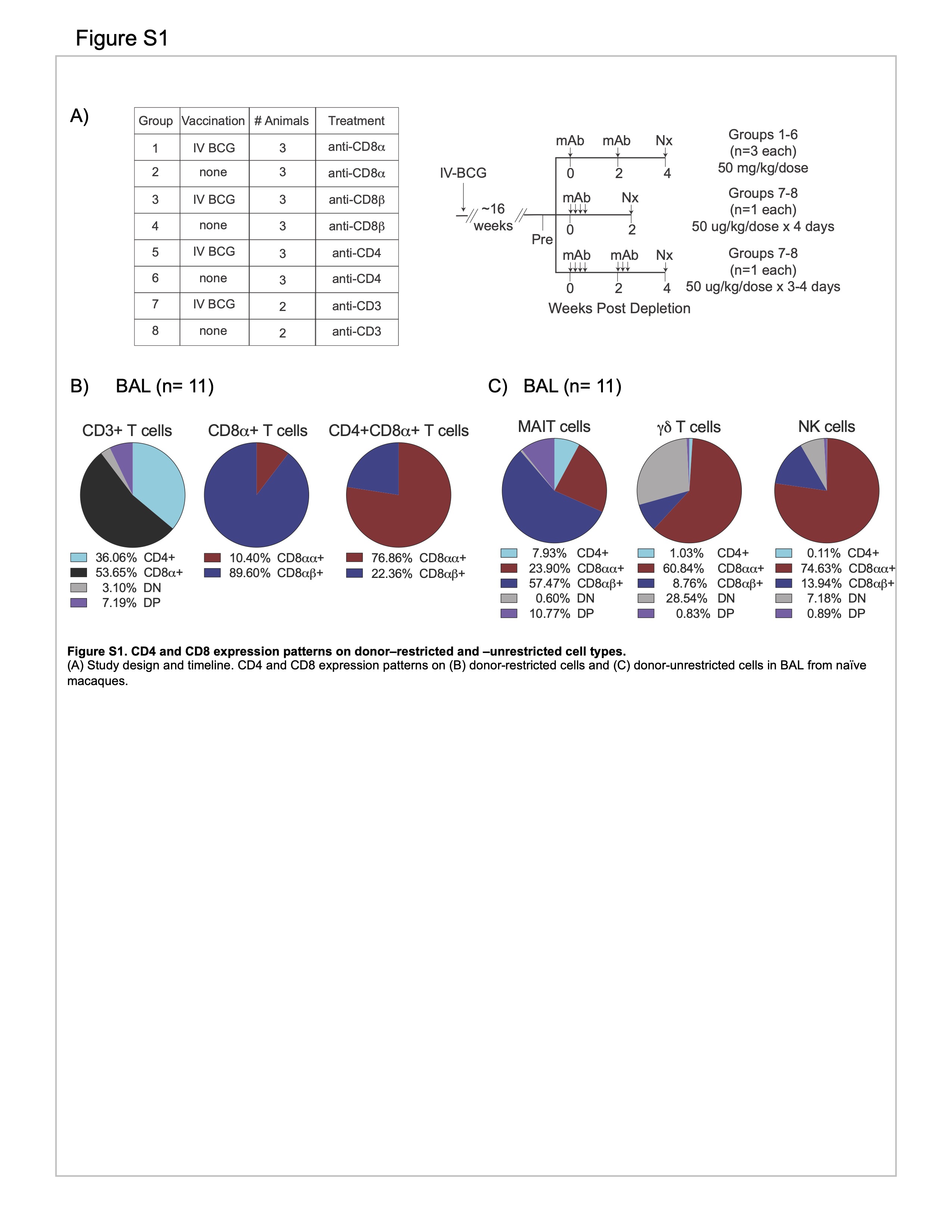

### Supplemental Figure 2

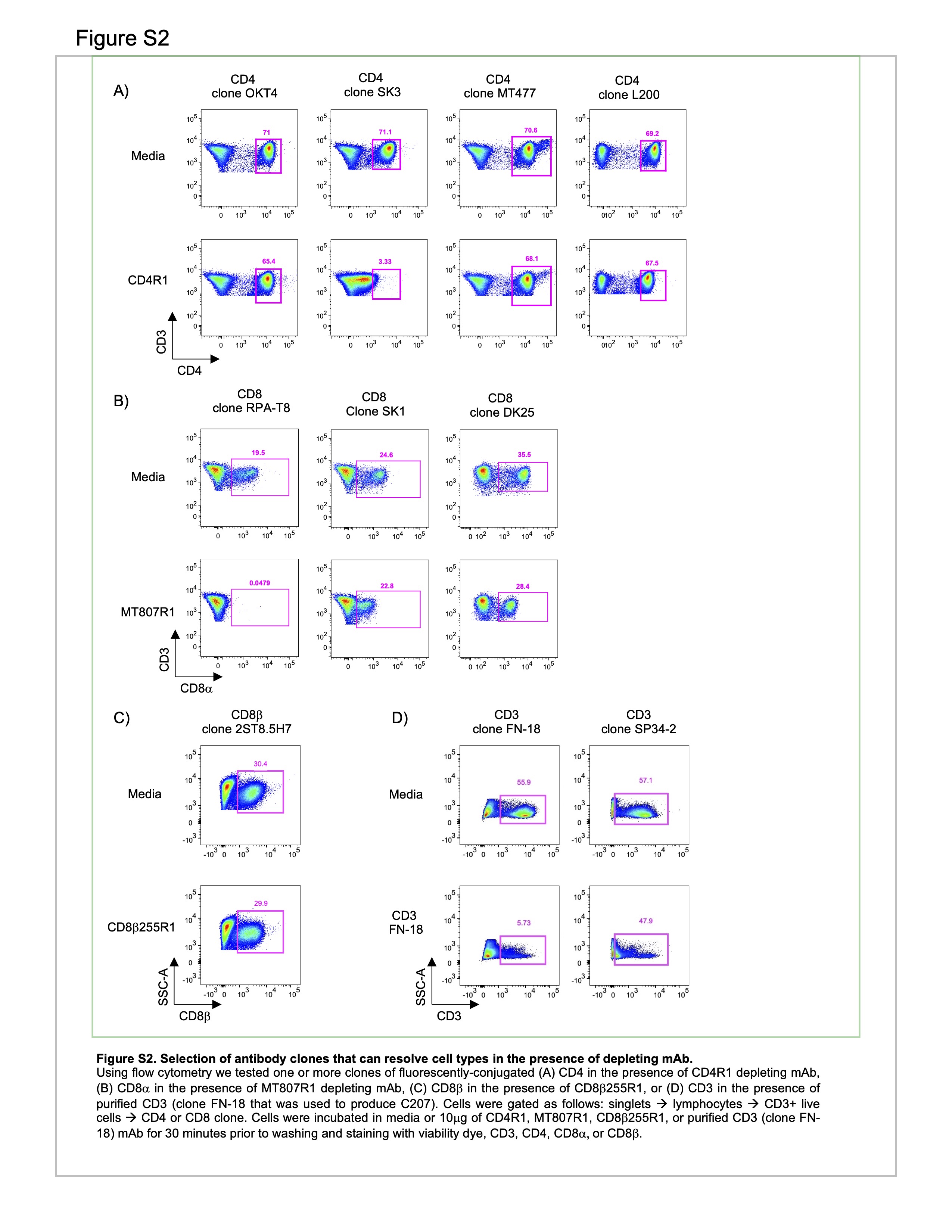

### Supplemental Figure 3

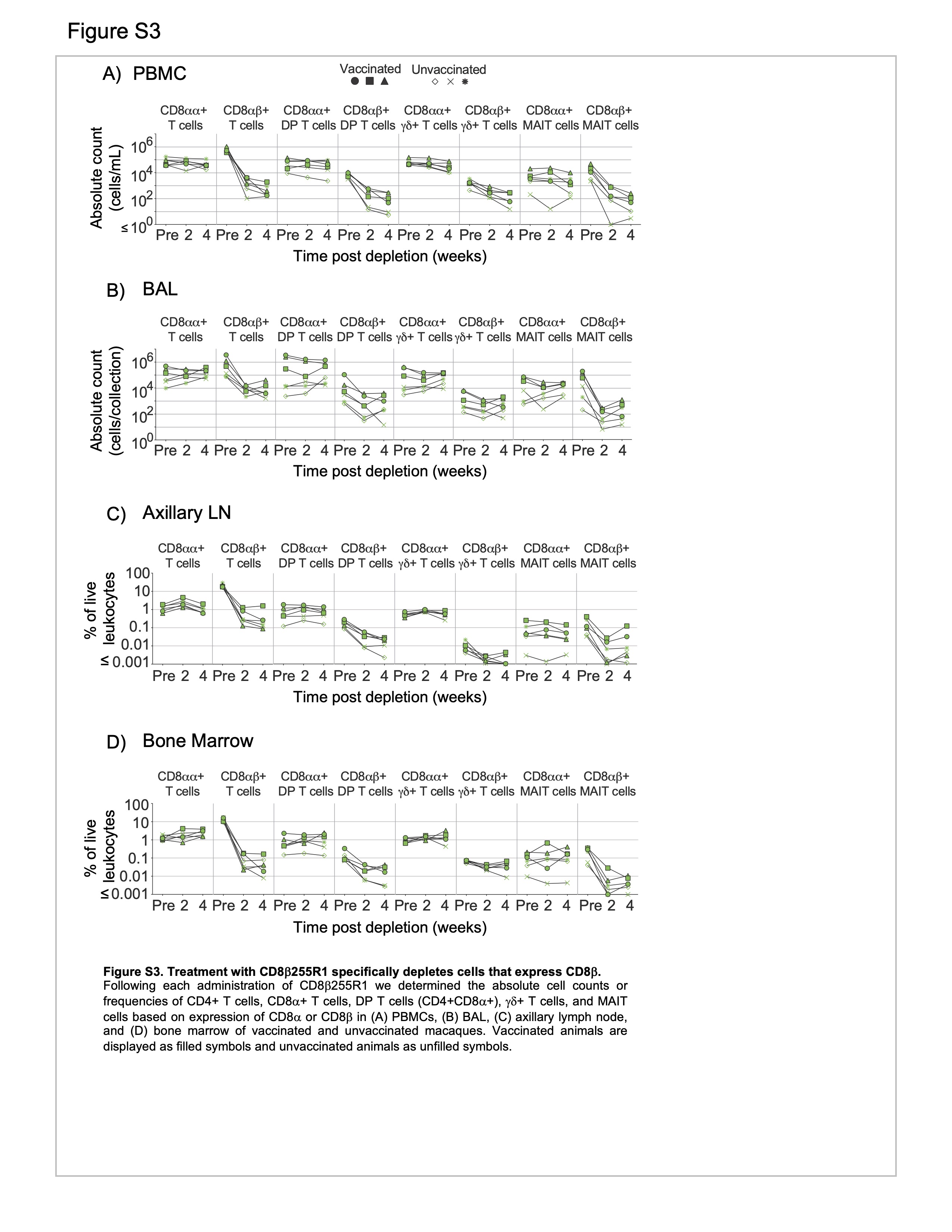

### Supplemental Figure 4

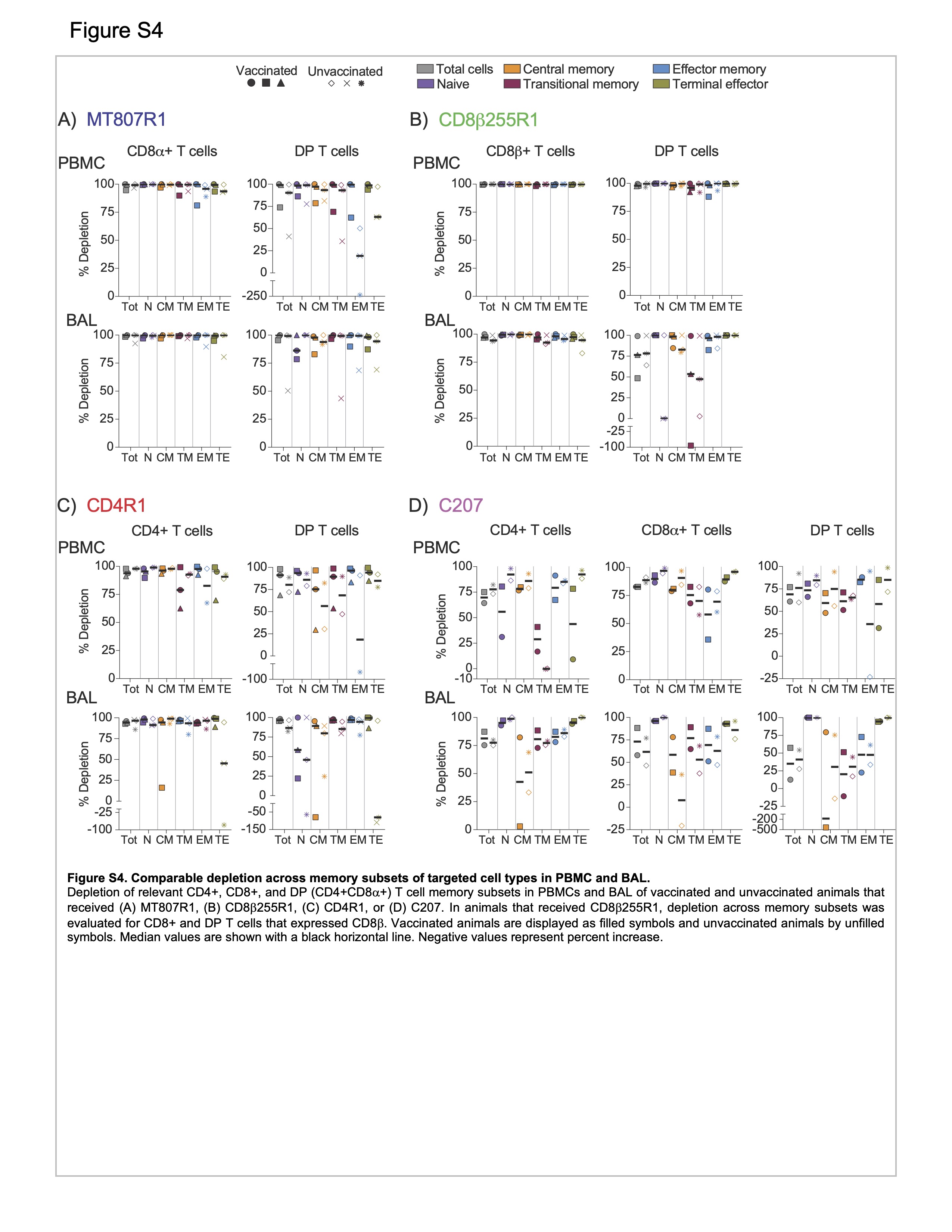

### Supplemental Figure 5

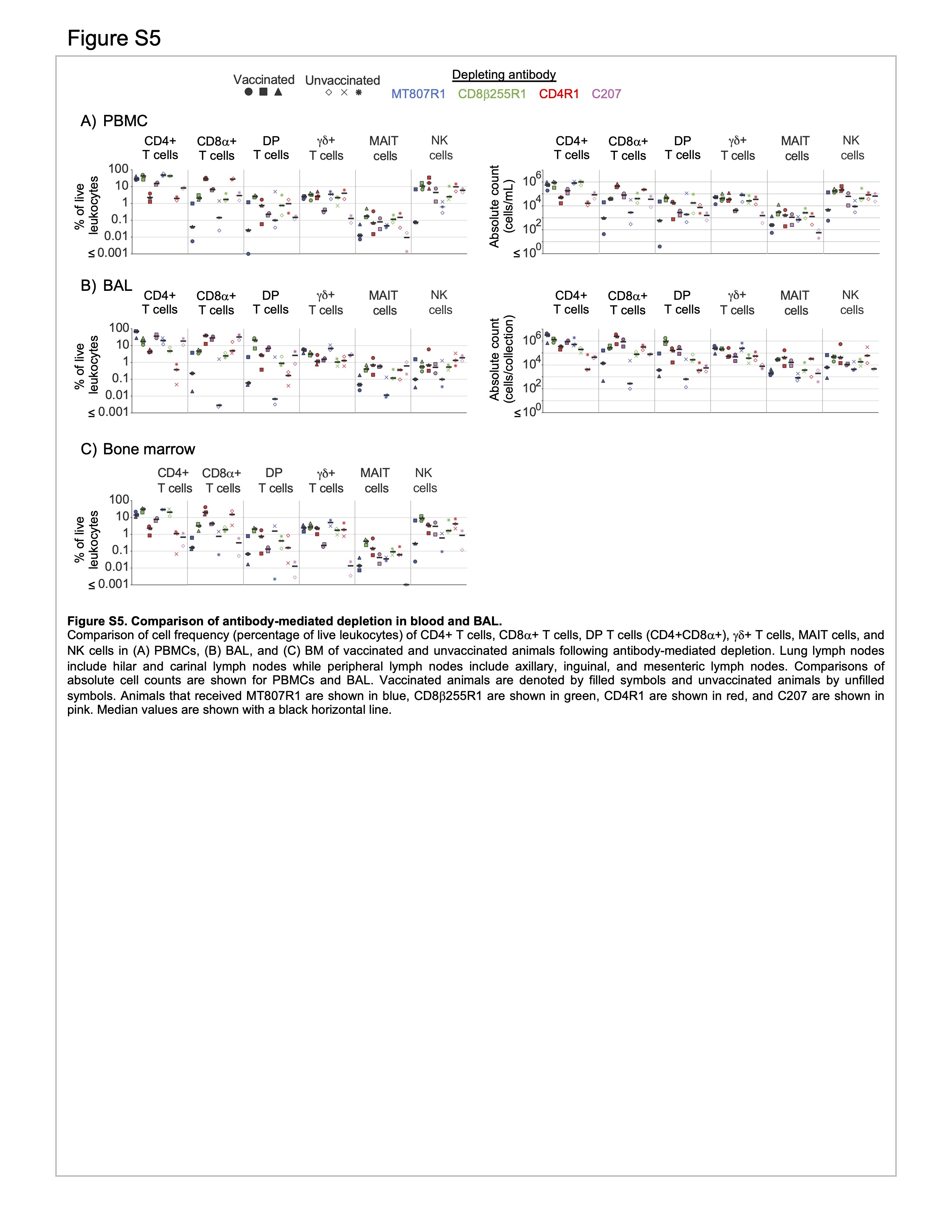

### Supplemental Figure 6

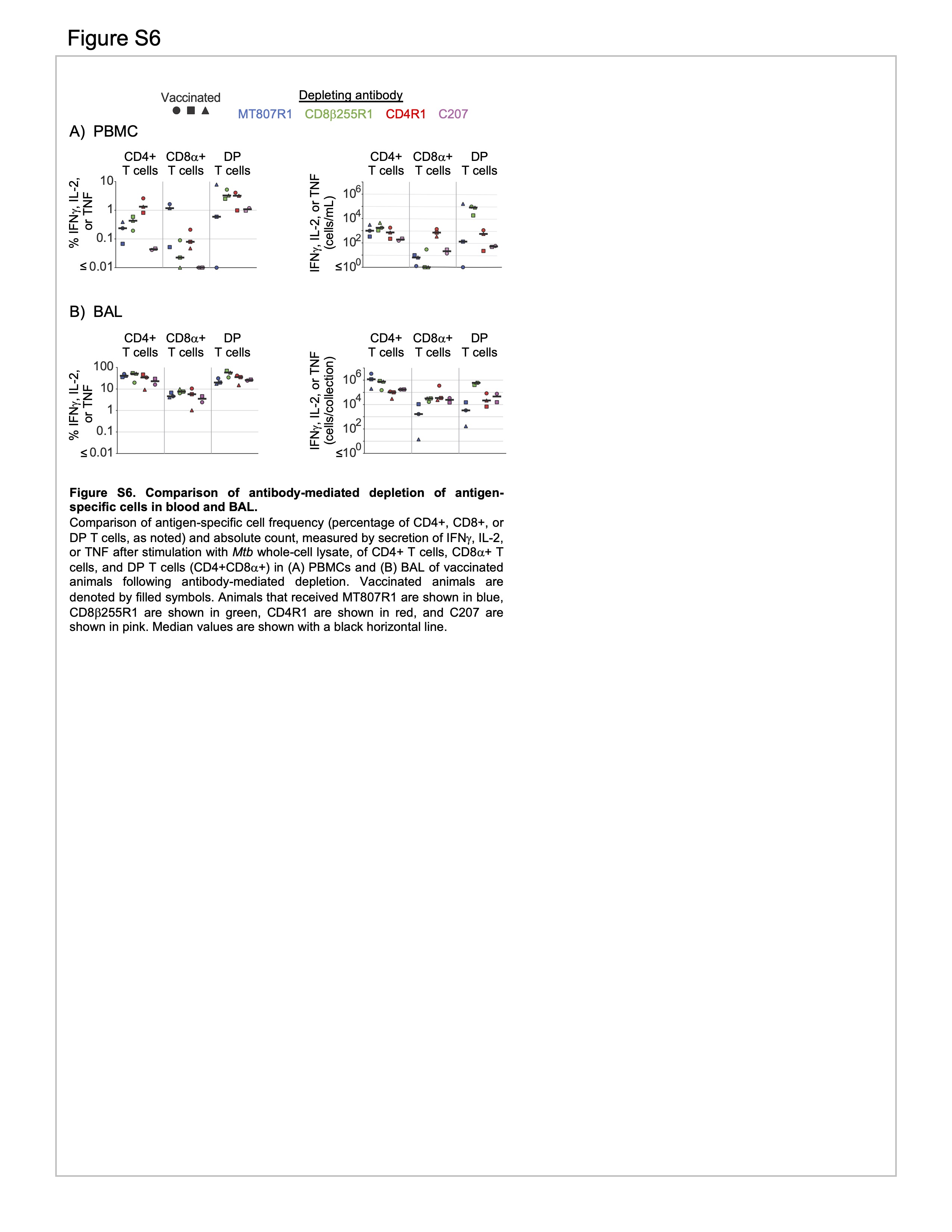
